# scMuffin: an R package for disentangling solid tumor heterogeneity from single-cell expression data

**DOI:** 10.1101/2022.06.01.494129

**Authors:** Valentina Nale, Alice Chiodi, Noemi Di Nanni, Ingrid Cifola, Marco Moscatelli, Cinzia Cocola, Matteo Gnocchi, Eleonora Piscitelli, Ada Sula, Ileana Zucchi, Rolland Reinbold, Luciano Milanesi, Alessandra Mezzelani, Paride Pelucchi, Ettore Mosca

## Abstract

**INTRODUCTION:** Single-cell (SC) gene expression analysis is crucial to dissect the complex cellular heterogeneity of solid tumors, which is one of the main obstacles for the development of effective cancer treatments. Such tumors typically contain a mixture of cells with aberrant genomic and transcriptomic profiles affecting specific sub-populations that might have a pivotal role in cancer progression, whose identification eludes bulk RNA-sequencing approaches. We present scMuffin, an R package that enables the characterization of cell identity in solid tumors on the basis of a various and complementary analyses on SC gene expression data.

**RESULTS:** scMuffin provides a series of functions to calculate qualitative and quantitative scores, such as: expression of marker sets for normal and tumor conditions, pathway activity, cell state trajectories, CNVs, transcriptional complexity and proliferation state. Thus, scMuffin facilitates the combination of various evidences that can be used to distinguish normal and tumoral cells, define cell identities, cluster cells in different ways, link genomic aberrations to phenotypes and identify subtle differences between cell subtypes or cell states. We analysed public SC expression datasets of human high-grade gliomas as a proof-of-concept to show the value of scMuffin and illustrate its user interface. Nevertheless, these analyses lead to interesting findings, which suggest that some chromosomal amplifications might underlie the invasive tumor phenotype and the presence of cells that possess tumor initiating cells characteristics.

**CONCLUSIONS:** The analyses offered by scMuffin and the results achieved in the case study show that our tool helps addressing the main challenges in the bioinformatics analysis of SC expression data from solid tumors.

## 1. Background

Single-cell (SC) gene expression analysis is crucial to dissect the complex cellular heterogeneity of solid tumors, which is one of the main obstacles for the development of effective cancer treatments (1). A relevant number of software tools has been developed in recent years in the field of SC data analysis (2), a fact that stresses the key opportunities and challenges in this field. A recent study has shown that the development of tools that address common tasks (e.g., clustering of similar cells) and ordering of cells (e.g., definition of cell trajectories) is decreasing, while a greater focus is being paid on data integration and classification (2). These observations reflect the growing availability, scale and complexity of SC datasets (2).

SC datasets of solid tumors are typical examples of complex datasets that present a series of computational challenges and whose analysis demands domain-specific and integrative approaches. In fact, solid tumors typically contain a mixture of cells with aberrant genomic and transcriptomic profiles affecting specific sub-populations that might play a pivotal role in cancer progression, whose identification eludes bulk RNA-sequencing approaches. The use of cell type-specific markers (when available) is limited, and the alterations of gene expression that mark cancer cells makes the use of markers for normal cells not completely adequate. Moreover, the molecular heterogeneity of cancer cells (due to both intra-tumor and inter-individual differences) poses intrinsic limits in the definition of such markers. In addition, solid tumor samples typically comprise cells from the surrounding tissue or infiltrating cells that need to be distinguished from tumor populations for an effective analysis. Another challenge is the identification of clinically relevant cell subtypes that may be rare in the tumor mass, such as cancer stem cells or drug resistant subclones: because of their relatively low number, these cells are typically clustered together with many others. Lastly, an intrinsic problem of many SC datasets is the sequencing depth limit at the SC level. These limitations bound the number of detectable genes to the few thousands of the highest expressed genes, which implies, for example, that some established markers may not be used for data analysis.

To address these challenges, we developed scMuffin, an R package that implements a series of complementary analyses aimed at shedding light on the complexity of solid tumor SC expression data, including: a fast and customizable gene set scoring system; gene sets from various sources, including pathways, cancer functional states and cell markers; cell cluster association analysis with quantitative (e.g., gene set scores) as well as categorical (e.g. mutation states, proliferation states) features; copy number variation (CNV) analysis; transcriptional complexity analysis; proliferation rate quantification; and gene set based multi-dataset analysis **(Figure 1)**. scMuffin facilitates the integrative analysis of these multiple features, thus allowing the identification of cell subtypes that elude more general clustering and classification approaches. We describe the key aspects of scMuffin implementation and then its user interface by means of a case study on a public SC expression dataset of human high-grade gliomas (HGG).

**Figure 1.**
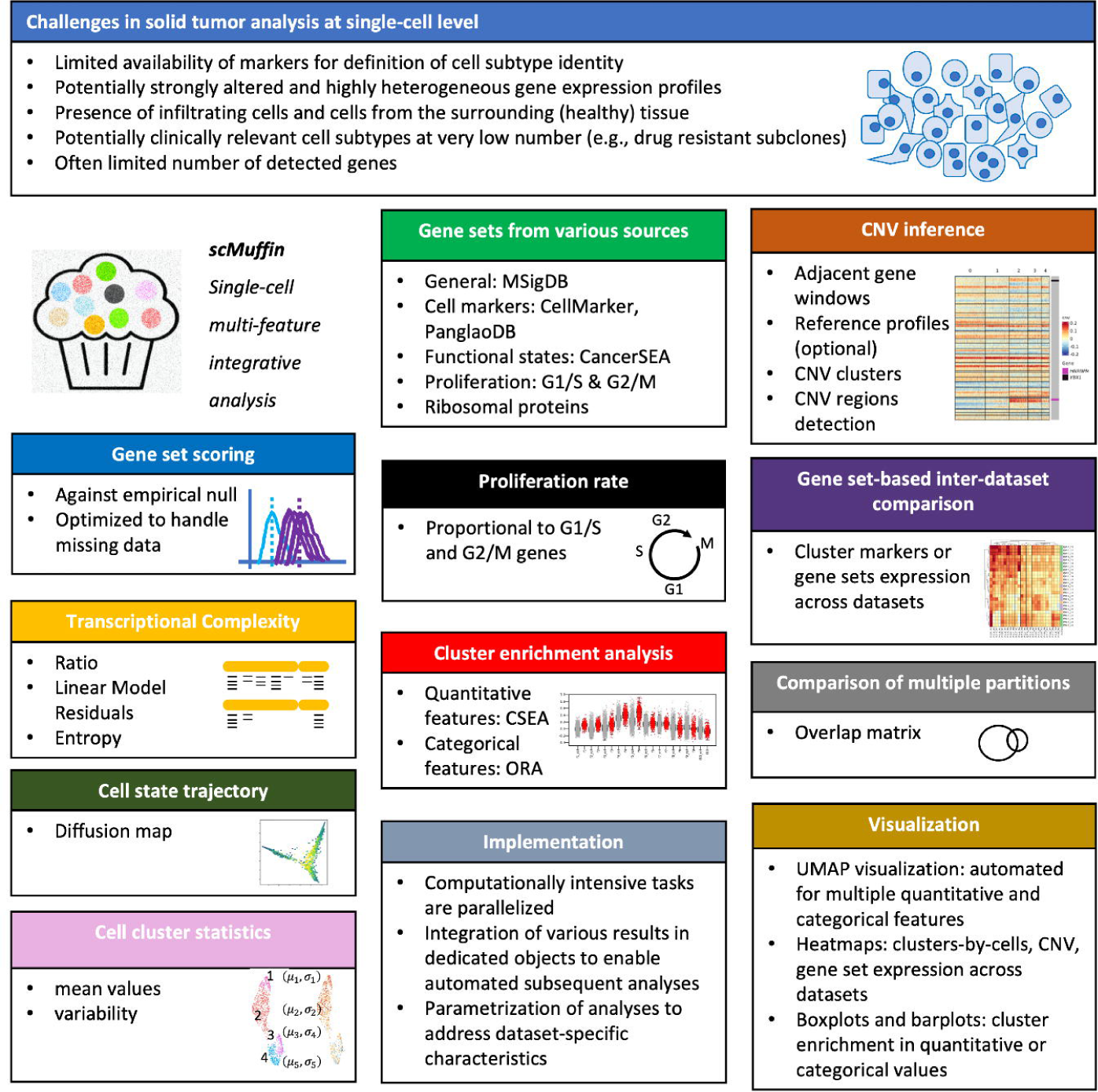
Overview of scMuffin package. scMuffin offers the possibility to perform several different analyses and data integration approaches to address the main challenges of SC gene expression analysis in solid tumors.

## 2. Implementation

scMuffin is implemented as an R package and provides a series of functions that allows the user to quantify various genetic and phenotypic characteristics of single cells, which can be combined to obtain insights on cell identity and function. The functions of scMuffin operate on a common data structure, the “scMuffinList”, by adding, changing or removing its elements. Computationally intensive tasks (in particular, gene set scoring and CNV inference) are parallelized. Package documentation includes R vignettes and function documentation, which are also available as “GitHub Pages” at the URL https://emosca-cnr.github.io/scMuffin. In this section, we describe package inputs and the main algorithms and definitions underlying the analyses offered by scMuffin.

### 2.1 Input

scMuffin is intended to be used downstream to general purpose tasks like quality control, normalization, cell clustering and dataset integration, for which there are dedicated tools, such as Seurat (3). scMuffin requires the genes-by-cells raw counts matrix, genes-by-cells normalized expression matrix and a partition of cells in clusters. Typically, these matrices are filtered during preprocessing steps to exclude low quality cells and genes that could negatively affect the analyses. However, the characterization of cells that can be achieved with scMuffin offers insights that can be used to (further) filter the dataset and/or to decide on which cells apply particular analyses (e.g. biomarker identification). In general, according to research questions and experimental design one may want to apply strong or mild filters before using scMuffin.

### 2.2 Quantification of gene set expression scores at cell and cluster levels

The quantification of gene set expression scores follows the approach described in (4,5), in which a gene set is scored on the basis of its average deviation from an empirical null. The approach is implemented enabling the user to tune various parameters (e.g., the minimum number of cells in which a gen set must be expressed, the number of bins of the null model and the possible removal of missing values) to better address the needs of a particular study in relation to the characteristics of the dataset under analysis. In particular, the code is implemented to handle missing values, which are one of the main issues in SC datasets. For example, this is important to quantify the expression of gene sets that are expressed just in a small subset of cells (and have null values elsewhere), or to account for the expression of different members of the same pathway in different cells.

The algorithm for gene set scoring is the following. Given a gene set *s*:

1. the genes occurring in the normalized genes-by-cells matrix are grouped into a series of bins according to their average expression across cells;
2. a number *k* of random gene sets 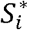are created, of the same size of *s*, tossing genes from the same bins of *s*, in order to match the distribution of gene expression of each 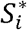with that of s;
3. the averages *m*_C_ and 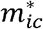are calculated, respectively, over the values of *s* and 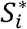in every cell *c*;
4. the expression score *Y*_*c*_ is calculated as the average difference between *m*_*c*_ and 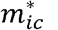

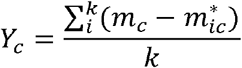
5. the difference between the medians of *m*_C_ and 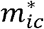 values in any cluster is used as the cluster-level score of *s*.

### 2.3 CNV estimation based on adjacent gene expression

CNV inference in scMuffin is based on the “adjacent gene windows” approach (4,6), which has been validated using both single nucleotide polymorphism arrays (6) and whole-exome sequencing (4). The approach is implemented in parallel and, like gene set scoring, offers various parameter tuning and data filtering possibilities, which allow the investigator to optimize the analysis to specific needs. The CNV profile of each cell is calculated as a moving average of scaled gene expression levels ordered by genomic location. scMuffin offers the possibility of subtracting a “normal” reference profile to highlight relative CNVs. Since a SC dataset derived from a sample of a solid tumor can either contain or not a proper set of physiological cells to be used as reference, this reference can be derived from cells that are part of the dataset under analysis (as described in (4)), or using external dataset (as described in (6)). The main steps are:

1. the reference cells are added to the genes-by-cells matrix (optional);
2. the expression of each gene is scaled subtracting its average (optional);
3. the gene expression matrix is ordered by chromosome and gene location;
4. in each chromosome *h*, the estimated copy number *V*_iC_ of cell *c* is calculated for all the ordered genes 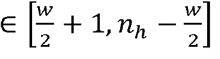

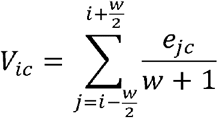

where *w* is an even number that defines the window size, that is, the number of genes located before and after gene *i* which contribute to the estimation of *V*_ic_, *n*_h_ is the number of genes in *h*, and e_*jc*_ is the gene expression value;
5. *V*_*ic*_ values are scaled subtracting their average in a cell (optional);
6. cells are clustered by their CNV profile;
7. the average CNV profile of the normal reference cells is subtracted from all the CNV profiles (optional).
8. the summary CNV score of a cell is calculated as:

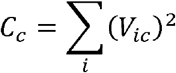

Cells are clustered based on their CNV profile using Seurat (3). A CNV region of a cluster is composed of the adjacent genes *i* that have absolute median *V*_ic_(within the cluster) higher than a given threshold. This threshold is by default defined as the standard deviation of *V*_ic_ values of all cells.

### 2.4 Transcriptional complexity

The transcriptional complexity (TC) is calculated using three different approaches. The TC-ratio 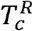of a cell is inferred based on the number of expressed genes (g_c_) over the number of total mapped reads (transcripts) (t_c_):

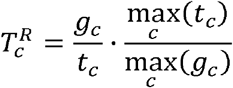

where g_c_ =#{r_ic ≥α} α_ is a threshold over the gene count r_ic_ and defines the gene *i* as “expressed”, and tc = Σ i r_*ic*_. Values of 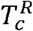higher (lower) than 1 indicate a higher (lower) number of expressed genes in relation to the total transcripts of the cell.

The TC-LMR (linear model residual) of a cell corresponds to its residual in the linear regression model between g_c_ and t_c_ :

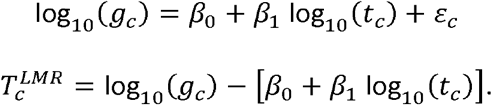

Positive (negative) residuals indicate higher (lower) number of expressed genes in relation to total transcripts.

The TC-H (entropy) is quantified as the transcriptional entropy of the cell (7):

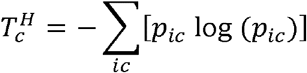

where 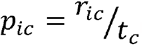 The higher the entropy the higher the number of expressed genes.

### 2.5 Proliferation rate

The proliferation rate P_*c*_ of a cell is quantified as the maximum between the two gene set scores *Y*_*c*_(S1)and *Y*_*c*_(S2), calculated on the gene sets *s*_1_ and *s*_2_ that contain, respectively, genes involved in G1/S and G2/M cell cycle phases:

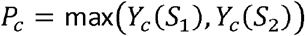

where *s*_1_ and *s*_2_ are defined as in Tirosh *et al*. (4).

### 2.6 Cluster enrichment analysis for quantitative and categorical features

The association between cell clusters and any cell-level values is performed through Cell Set Enrichment Analysis (CSEA) and Over Representation Analysis (ORA) for, respectively, quantitative and categorical values. Importantly, these two types of analysis are implemented in parallel, which is particularly important for CSEA that uses permutations to build an empirical null distribution.

CSEA is the application of Gene Set Enrichment Analysis (GSEA) (8) to cells and cell sets in place of, respectively, genes and gene sets: a ranked cell list is used instead of a ranked gene list, and cell sets (cell clusters) are tested instead of gene sets. In addition, the code is implemented to handle missing values. Therefore, scMuffin tests whether the cells assigned to a cluster are located at the top (bottom) of a ranked list of cells.

The assessment of cluster enrichment in a particular value of a categorical feature is computed using the over-representation analysis (ORA) approach (9), which is based on the hypergeometric test. Here, this test assesses whether the occurrence of a particular value between the cells of a cluster in relation to all other clusters is higher than expected in a hypergeometric experiment.

### 2.7 Overlap between partitions of cells

Cell partitions resulting from different clustering analyses are compared by calculating the overlap coefficients among all-pairs of clusters. Given two partitions *A* and *B*, defined as sets of cell

clusters *A*={*a*_*i*_*}* andB={*b*_*j*_*}* the similarity between the cell clusters *a*_i_ and *b*_J_ is calculated as:

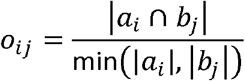

## 3. Results and Discussion

In this section, we present the user interface **(Table 1)**, taking advantage of the results obtained on the SC dataset generated by Yuan *et al*. (10) from human high-grade glioma (HGG) samples (available on the Gene Expression Omnibus (GEO) repository (11) with identifier GSE103224, see Supplementary Methods in Additional File 1).

**Table 1.** Main tasks and corresponding functions in scMuffin.

| Task | Description | User interface |
| --- | --- | --- |
| Create the scMuffinList | Create the scMuffinList from counts and normalized expression data | `create_scMuffinList()` |
| Add a partition | Add a partition of cell (clusters) | `add_partition()` |
| Gene set expression scoring | Average gene set expression deviation from matched empirical reference;<br>provided gene sets from CellMarker (12), PanglaoDB (13), CancerSEA (14) and MSigDB (8) | `prepare_gsls()`<br>`calculate_gs_scores()`<br>`calculate_gs_scores_in_clusters()` |
| CNV | Estimation of CNVs by means of the “moving window” approach, that is, considering the expression of adjacent genes;<br>calculation of CNV deviation from a normal reference profile;<br>processing of normal tissue-specific expression data from GTEx;<br>detection of CNV regions | `CNV_analysis()`<br>`process_GTEx_gene_reads()` |
| Transcriptional complexity | Number of expressed genes in relation to the total reads, calculated using a ratio, linear model residuals and entropy. | `transcr_compl()` |
| Proliferation | Maximum between G1/S and G2/M gene set scores | `proliferation_analysis()` |
| Cell state trajectory | Diffusion map computation | `diff_map()` |
| Cell cluster annotation | Assessment of cluster enrichment for quantitative and categorical features | `assess_cluster_enrichment()` |
| Comparison of datasets | Assessment of cluster markers or gene set expression across multiple datasets | `inter_dataset_comparison()` |
| Visualization | Automated UMAP visualizations for multiple features;<br>Heatmaps: gene set scores, CNV, overlap between clusters;<br>boxplots and barplots of cluster enrichment | `boxplot_cluster()`<br>`dotplot_cluster()`<br>`heatmap_CNV()`<br>`plot_profile_CNV()`<br>`plot_umap_colored_features()`<br>`plot_heatmap_features_by_clusters()` |

### 3.1 Gene set scoring

scMuffin provides functions to set up one or more gene set collections and perform SC-level estimation of gene set expression scores in relation to an empirical null model (see Implementation section). This can be applied to any gene set and can therefore be used to estimate several different cell phenotypes, like pathway activities or marker set expression.

The function `prepare_gsls()` allows the user to collect gene sets of cell types, pathways, cancer functional states, as well as other collections of gene sets (e.g. positional gene sets, hallmarks) from CellMarker (12), PanglaoDB (13), CancerSEA (14) and MSigDB (8) databases.

Unlike many existing tools that are used to perform marker-based cell annotation (15), the availability of these gene sets within scMuffin package spares the user the effort of data collection and harmonization. The function, which also accepts any user-given gene sets, applies a series of criteria (e.g., minimum and maximum number of genes in a gene set) to filter the chosen gene sets.

The cell-level expression scores for these gene sets can be calculated using `calculate_gs_scores()`. For instance, the following code shows how to quantify the activity of “Cancer functional states” from CancerSEA (14) at cell and cell cluster level:

~~~
gsc <-prepare_gsls(gs_sources = “CancerSEA”, scMuffinList = scMuffinList)
scMuffinList <-calculate_gs_scores(scMuffinList = scMuffinList, gs_list = gsc$CancerSEA)
scMuffinList <-calculate_gs_scores_in_clusters(scMuffinList = scMuffinList, partition_id = “global_expr”)
~~~

where “scMuffinList” is the data structure that contains expression data and cell clusters (“global_expr”).

Cluster-level gene sets scores can be visualized as a heatmap to obtain a useful summary visualization that can provide insights for cluster annotation. This is accomplished by means of the function `plot_heatmap_features_by_clusters()`, which relies on the powerful ComplexHeatmap R package (16)). For example, the analysis of the CancerSEA functional states in the HGG sample PJ016 showed that the two groups of cell clusters that are spatially separated in the UMAP visualization, that is clusters {0, 6, 8, 9} and {1, 2, 3, 4, 5, 7} **(Figure 2a)**, reflect distinct functional states **(Figure 2b)**. Cell-level gene set scores can be visualized over the UMAP by means of `plot_umap_colored_features()` **(Figure 2c)**.

**Figure 2.**
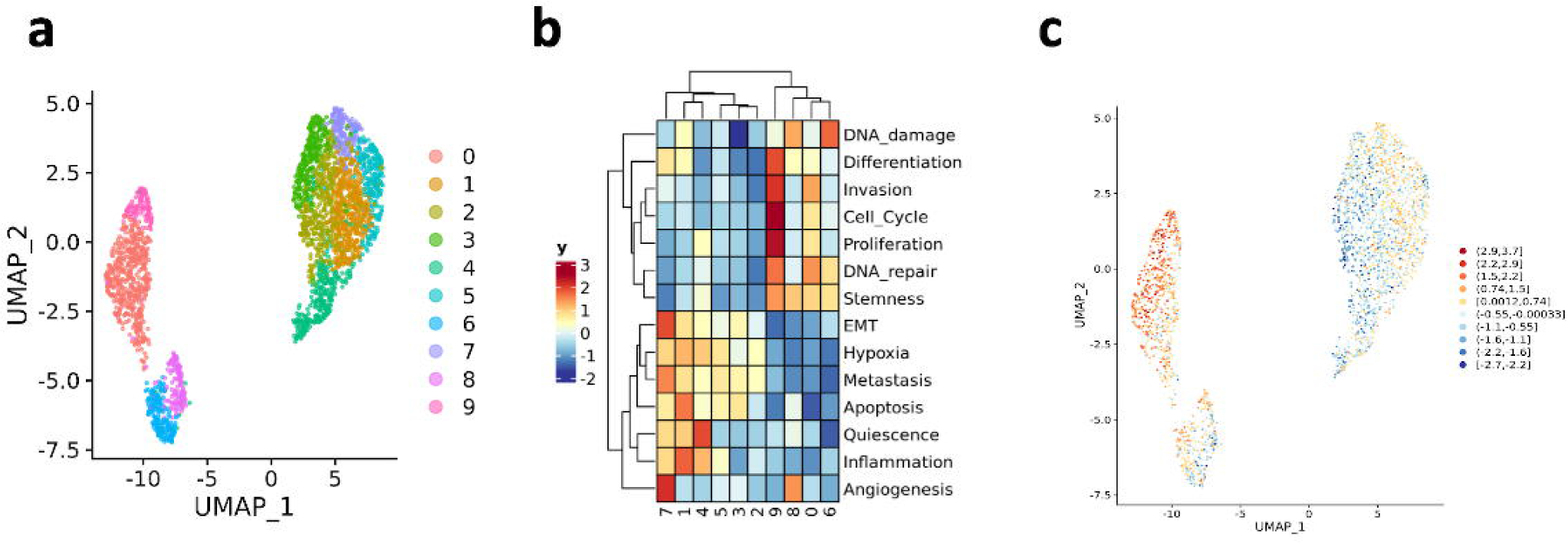
Quantification of CancerSEA functional states in the HGG sample PJ016. **a)** UMAP visualization where cells are coloured by expression clusters. **b)** Cluster-level expression scores of all the CancerSEA functional states. c) UMAP visualization where cells are colored by “CSEA_Invasion” gene set score.

### 3.2 CNV estimation and association with CancerSEA functional states

CNV inference from SC transcriptomics in cancer provides a means to assess the presence of relevant genomic aberrations (duplications and deletions) based on the expression of adjacent genes. This knowledge offers crucial clues to address the difficult task of distinguishing normal from malignant cells and provides quantitative information to reconstruct the tumor clonal substructure. Moreover, CNV patterns allow the investigator to hypothesize link between genomic alterations and cell phenotypes. As a proof-of-concept, we describe CNV inference in scMuffin with and without a reference expression profile, and the association of CNV clusters with CancerSEA functional states.

CNV inference is performed by means of the function `CNV_analysis()`:

scMuffinList <-CNV_analysis(scMuffinList, reference = GTEx_mean, center_genes = TRUE)

where GTEx_mean contains a reference expression profile, which can be obtained by means of the function `process_GTEx_gene_reads()` starting from data available at The Genotype-Tissue Expression (GTEx) portal (17). The resulting scMuffinList contains the matrix with CNV values for each genomic regions in every cell, cell clusters by CNV, detected CNV regions and mapping information between gene locations and CNV regions.

Heatmaps that show CNV profiles of every cell, clustered by similarity, can be generated by means of `plot_heatmap_CNV()`. In the considered dataset (PJ030), we observed that the reference profile (normal brain samples from GTEx) is included into cluster 3, while clusters 0, 1 and 2 show large aneuploidies, some of which are typical of HGG, like the amplification of chromosome 7 as reported by dataset authors (10) **(Table 2, Figure 3a-b)**. The median CNV profile of every cell cluster can be visualized by means of `plot_profile_CNV()` **(Figure 3c)** while boxplots of the distribution of summary CNV score by clusters can be visualized through `boxplot_points()` **(Figure 3d)**.

**Table 2.**
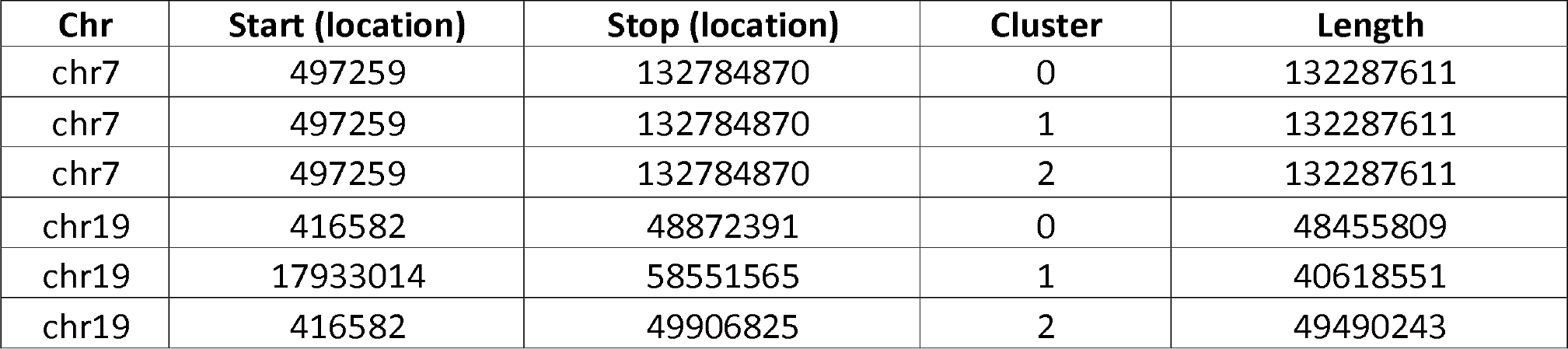
CNV regions detected in chromosomes 1 and 7 of sample PJ030.

| Chr | Start (location) | Stop (location) | Cluster | Length |
| --- | --- | --- | --- | --- |
| chr7 | 497259 | 132784870 | 0 | 132287611 |
| chr7 | 497259 | 132784870 | 1 | 132287611 |
| chr7 | 497259 | 132784870 | 2 | 132287611 |
| chr19 | 416582 | 48872391 | 0 | 48455809 |
| chr19 | 17933014 | 58551565 | 1 | 40618551 |
| chr19 | 416582 | 49906825 | 2 | 49490243 |

**Figure 3.**
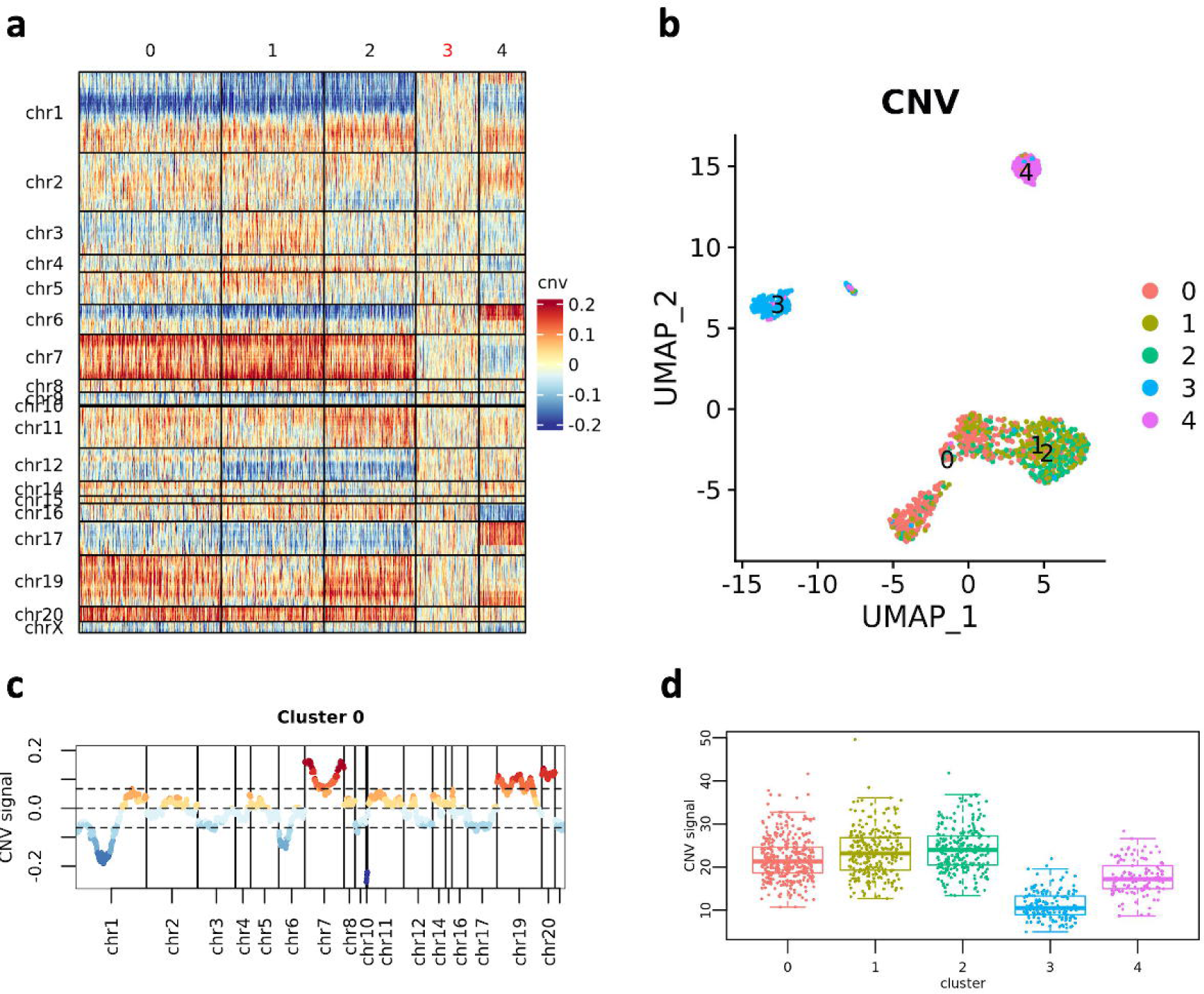
CNV analysis (PJ030) **a)** CNV heatmap where cells (columns) are grouped into “CNV clusters”; the cluster of cells that include the reference profile is highlighted through a red label. **b)** UMAP visualizations where cells are colored by CNV clusters. **c)** Median CNV value of a cluster 0. d) Distribution of the summary CNV score in clusters.

The analysis of another dataset (PJ016, without reference profile) showed two groups of clusters ({0, 1, 4} and {2, 3}) that map to separated regions of the UMAP visualization **(Figure 4a-b)**. Interestingly, clusters 2 and 3 are marked by peculiar amplifications in chromosome arms 1p and 19p. These “CNV clusters” correspond to expression clusters {0, 6, 8, 9} **(Figure 4c)**. The similarity of multiple cell partitions can be quantified using the function `overlap_matrix()`.

**Figure 4.**
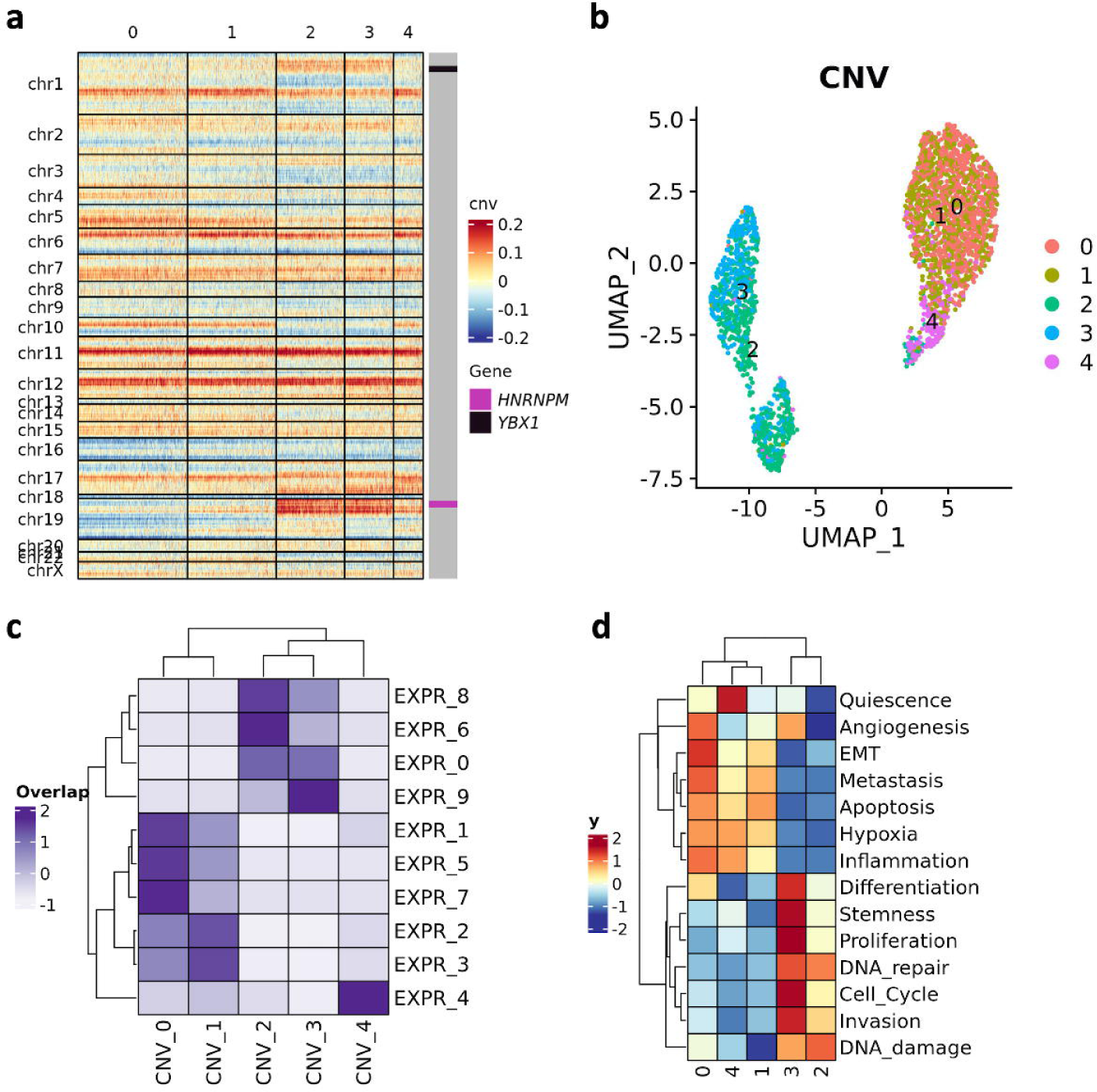
CNV analysis and CancerSEA functional states (PJ016) **a)** CNV heatmap where cells (columns) are grouped into “CNV clusters”. **b)** UMAP visualizations where cells are colored by CNV clusters. **c)** Overlap indices between expression clusters (prefix “EXPR_”) and CNV clusters (prefix “CNV_”). d) Heatmap of CancerSEA functional state expression score in CNV clusters.

An example of integrative analysis enabled by scMuffin is the functional assessment of CNV clusters. We quantified the expression scores of CancerSEA functional states throughout the CNV clusters of sample PJ016. As expected, the two aforementioned groups of CNV clusters {0, 1, 4} and {2, 3} are characterized by different functional states **(Figure 4d)**. This finding suggests that the peculiar amplifications of chromosomes 1 and 19 that characterize these clusters might underlie two phenotypes. A suggestive evidence of such hypothesis, as regards the CancerSEA functional state “Invasion”, is the location of the two Invasion markers *YBX1* (Y-Box Binding Protein 1) and *HNRNPM* (Heterogeneous Nuclear Ribonucleoprotein M) within the amplified regions of chromosomes 1p and 19p **(Figure 4c)**. *YBX1* is a DNA/RNA-binding protein and transcription factor which plays a central role in coordinating tumor invasion in glioblastoma (18). *HNRNPM* belongs to a family of spliceosome auxiliary factors and is involved in the regulation of splicing; the upregulation of these factors results in tumor-associated aberrant splicing, which promotes glioma progression and malignancy (19,20). In particular, HNRNPM was identified as an interactor of the DNA/RNA binding protein SON, which drives oncogenic RNA splicing in glioblastoma (21). While it is beyond the scope of this article to validate this hypothesis, these findings clearly highlight the usefulness of the integrative analysis of CNVs and CancerSEA functional states provided by scMuffin.

### 3.3 Cluster enrichment analysis: association of clusters and features

scMuffin contains functions for assessing the association between cell clusters and quantitative as well as categorical features, by means of CSEA and ORA, respectively. The function `assess_cluster_enrichment()` performs CSEA and ORA accordingly to the feature type under consideration. For example, the following instruction runs CSEA over the gene set scores, considering the partition “global_expr”:

scMuffinList <-assess_cluster_enrichment(scMuffinList, feature_name = “gene_set_scoring”, partition_id = “global_expr”)

The results of CSEA and ORA include several quantities, such as the normalized enrichment score (NES), the enrichment ratio (ER) and the false discovery rate (FDR) **(Tables 3-4)**. These results can be visualized as boxplots (quantitative features) and barplots (categorical features). For instance, the cluster 0 of sample PJ016 is particularly enriched – as expected – in “Invasion” markers **(Table 3, Figure 5a)** and cells assigned to G2/M or S phases **(Table 4, Figure 5b)**. The function `extract_cluster_enrichment_table()` provides a means to obtain tables of enrichment results across clusters, which can be visualized as heatmaps **(Figure 5c)**, while `extract_cluster_enrichment_tags()` defines the most significant features as cluster labels that are therefore available for annotating UMAPs **(Figure 5d)**.

**Table 3.**
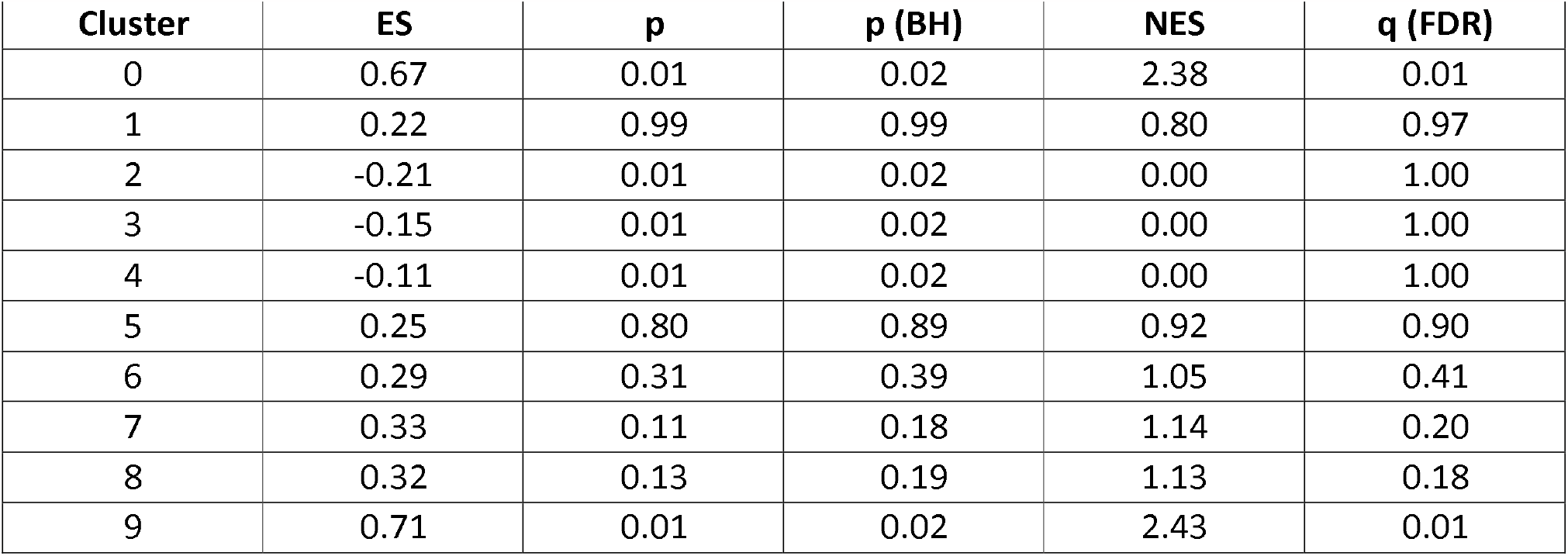
The output of CSEA analysis for a quantitative feature. This result is relative to the CancerSEA Invasion gene set on dataset PJ016. ES = Enrichment Score; p = CSEA p-value; p (BH) = p value after Bonferroni-Hochberg correction; NES = Normalized Enrichment Score; q (FDR) = q value of FDR.

| Cluster | ES | p | p (BH) | NES | q (FDR) |
| --- | --- | --- | --- | --- | --- |
| 0 | 0.67 | 0.01 | 0.02 | 2.38 | 0.01 |
| 1 | 0.22 | 0.99 | 0.99 | 0.80 | 0.97 |
| 2 | -0.21 | 0.01 | 0.02 | 0.00 | 1.00 |
| 3 | -0.15 | 0.01 | 0.02 | 0.00 | 1.00 |
| 4 | -0.11 | 0.01 | 0.02 | 0.00 | 1.00 |
| 5 | 0.25 | 0.80 | 0.89 | 0.92 | 0.90 |
| 6 | 0.29 | 0.31 | 0.39 | 1.05 | 0.41 |
| 7 | 0.33 | 0.11 | 0.18 | 1.14 | 0.20 |
| 8 | 0.32 | 0.13 | 0.19 | 1.13 | 0.18 |
| 9 | 0.71 | 0.01 | 0.02 | 2.43 | 0.01 |

**Table 4.**
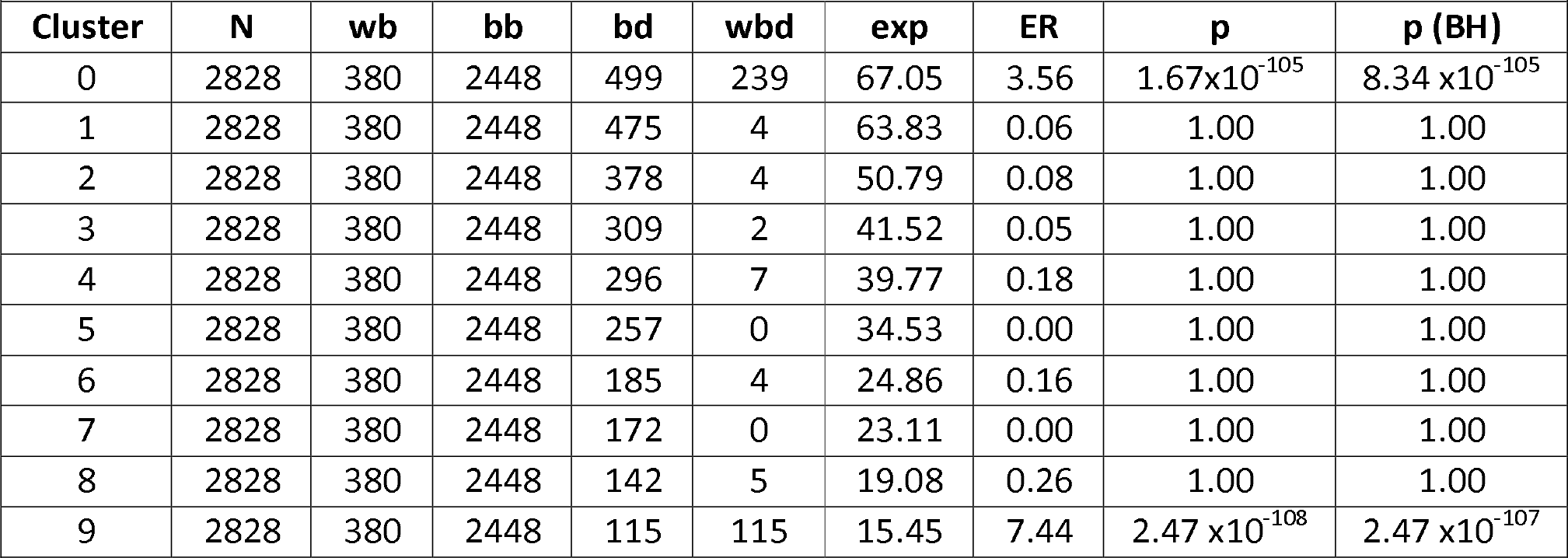
The output of an ORA analysis for a categorical feature. This result is relative to the cell cycle phase “G2M” on dataset PJ016. N=number of cells; wb = number G2M cells; bb = N-wb; wbd = number of G2M cells in the cluster; exp = expected number of G2M cells in the cluster; ER = enrichment ratio; p = hypergeometric p; p (BH) = p value after Bonferroni-Hochberg correction.

| Cluster | N | wb | bb | bd | wbd | exp | ER | p | p (BH) |
| --- | --- | --- | --- | --- | --- | --- | --- | --- | --- |
| 0 | 2828 | 380 | 2448 | 499 | 239 | 67.05 | 3.56 | $1.67 \times 10^{-105}$ | $8.34 \times 10^{-105}$ |
| 1 | 2828 | 380 | 2448 | 475 | 4 | 63.83 | 0.06 | 1.00 | 1.00 |
| 2 | 2828 | 380 | 2448 | 378 | 4 | 50.79 | 0.08 | 1.00 | 1.00 |
| 3 | 2828 | 380 | 2448 | 309 | 2 | 41.52 | 0.05 | 1.00 | 1.00 |
| 4 | 2828 | 380 | 2448 | 296 | 7 | 39.77 | 0.18 | 1.00 | 1.00 |
| 5 | 2828 | 380 | 2448 | 257 | 0 | 34.53 | 0.00 | 1.00 | 1.00 |
| 6 | 2828 | 380 | 2448 | 185 | 4 | 24.86 | 0.16 | 1.00 | 1.00 |
| 7 | 2828 | 380 | 2448 | 172 | 0 | 23.11 | 0.00 | 1.00 | 1.00 |
| 8 | 2828 | 380 | 2448 | 142 | 5 | 19.08 | 0.26 | 1.00 | 1.00 |
| 9 | 2828 | 380 | 2448 | 115 | 115 | 15.45 | 7.44 | $2.47 \times 10^{-108}$ | $2.47 \times 10^{-107}$ |

**Figure 5.**
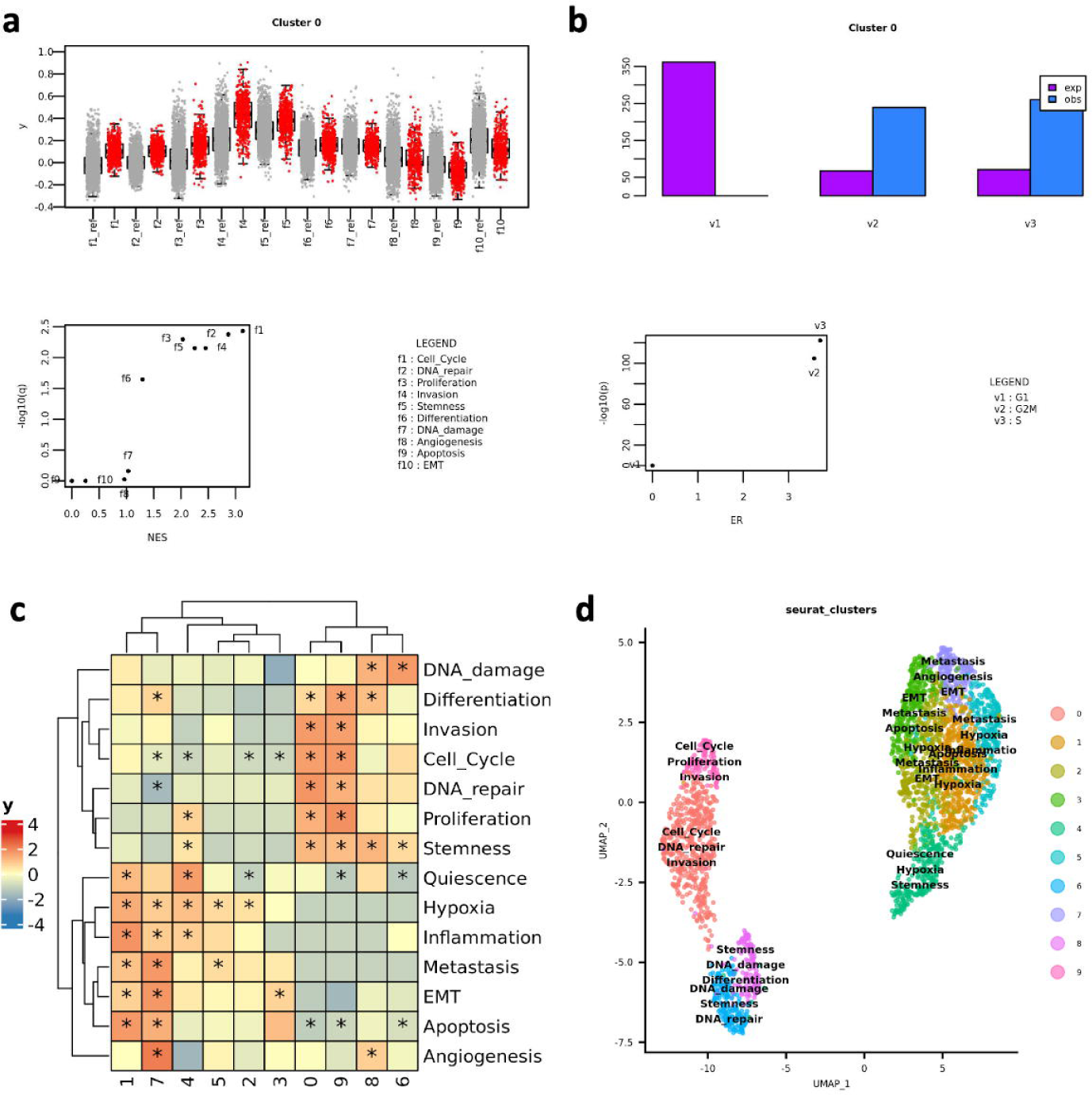
Cluster enrichment analysis (PJ016) **a)** Output of `boxplot_cluster()` function for a quantitative feature – CancerSEA functional states – in cluster 0: distribution of gene set scores values (red) and corresponfing null models (gray, suffix “_ref”) (top); Normalized Enrichment Score (NES) and FDR q-value (bottom). **b)** Output of `barplot_cluster()` function for a categorical feature – cell cycle phase – in cluster 0: expected (“exp”) and observed (“obs”) occurrences of every possible category (top); enrichment ratio (ER) and hypergeometric p-value (bottom). **c)** Heatmap of NES values for CancerSEA functional states in expression clusters; asterisks indicate FDR q-value < 0.01. **d)** UMAP visualization where clusters are labelled with the top three most significant CancerSEA functional states.

### 3.4 Transcriptional complexity, proliferation rate and cell state trajectories

The number of expressed genes and proliferation rate are two relevant pieces of information for the characterization of cell type and cell state in cancer.

Undifferentiated stem or progenitor cells show a high number of expressed genes (open chromatin state) compared to differentiated cell types. In cancer context, a high number of expressed genes might suggest tumor initiating cells (TICs), might indicate de-differentiation processes of tumor progression (22,23). For instance, in a recent study on glioblastoma, chromatin accessibility was associated to a specific subpopulation of putative tumor-initiating Cancer Stem

Cells (CSCs) with invasive phenotype and low survival prediction (24). The transcriptional complexity of a cell can be calculated as follows:

scMuffinList <-transcr_compl(scMuffinList, min_counts=3, min_cells=10, min_genes=500)

where min_counts is the threshold above which a gene is considered expressed, min_cells is the minimum number of cells in which a gene must be expressed and min_genes is the minimum number of genes that a cell must express. In SC datasets, transcriptional complexity quantification is complicated by the presence of a linear relationship between number of expressed genes and total transcripts detected, where biological variability is entangled with sequencing deepness. scMuffin provides three measures of transcriptional complexity, which show three different – but complementary – relations with the number of total transcripts (*t*): the ratio (TC-ratio) prioritizes cells with a relatively low *t* **(Figure 6a);** the linear model residuals (TC-LMR) are independent from *t* **(Figure 6b)**; lastly, the transcriptional entropy **(TC-H)** (7) highlights genes with relatively high number of *t* **(Figure 6c)**. All three measures may provide useful insights for the identification of particular cell types or states. For example, TC-ratio could be used when one is not interested in cell with a relatively abundant RNA production, like can be the case of quiescent cells with stem-like properties.

**Figure 6.**
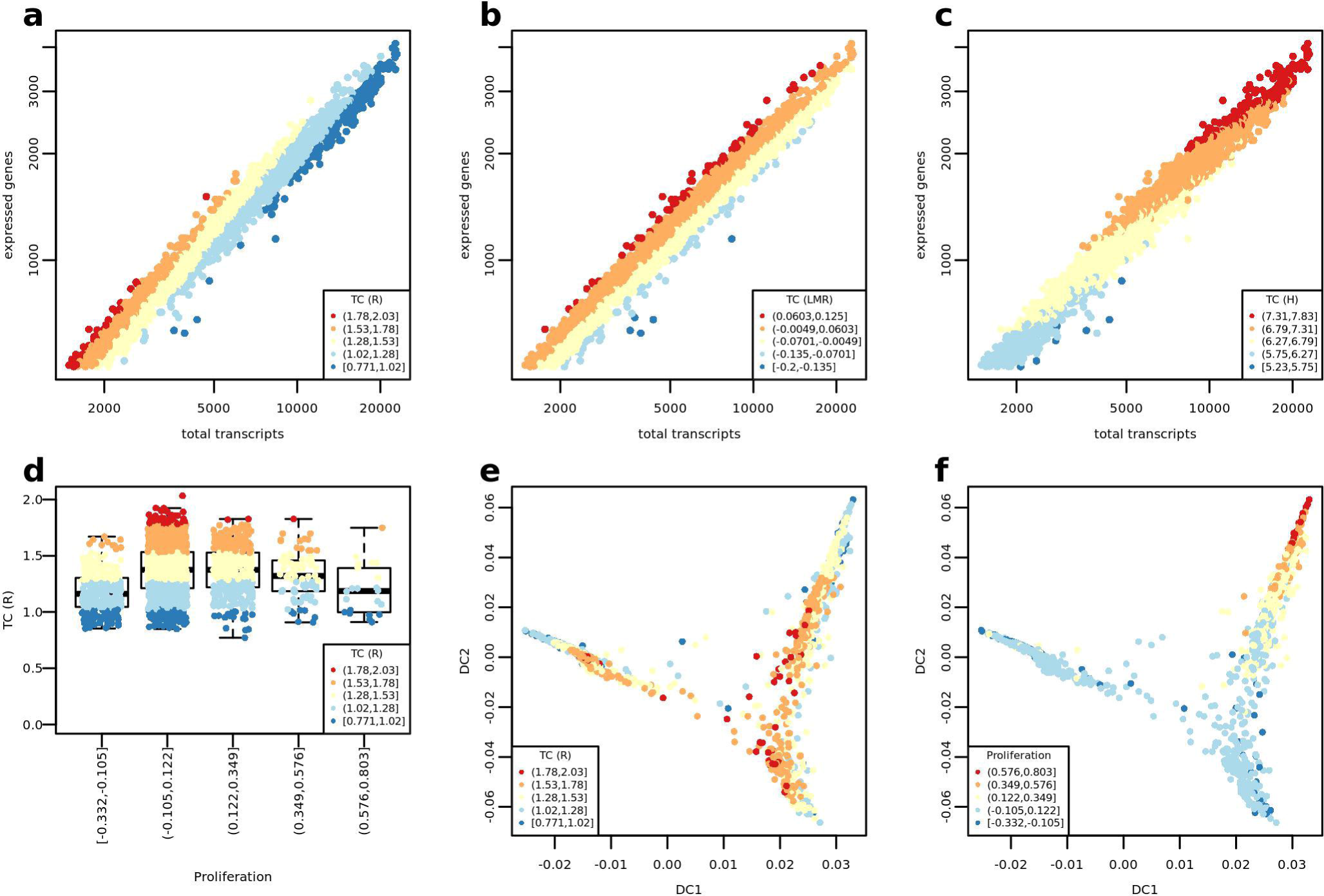
Transcriptional complexity, proliferation rate and cell state trajectories (PJ016) **a)**. Transcriptional Complexity Ratio (TC-R) of cells. **b)** Transcriptional Complexity Linear Model Residuals (TC-LMR) of cells. **c)** Transcriptional Complexity Entropy (TC-H) of cells. d) Distribution of TC-R values split by cell proliferation score. **e-f)** Distribution of cells according to the first two “diffusion components” (DC).

**Figure 7.**
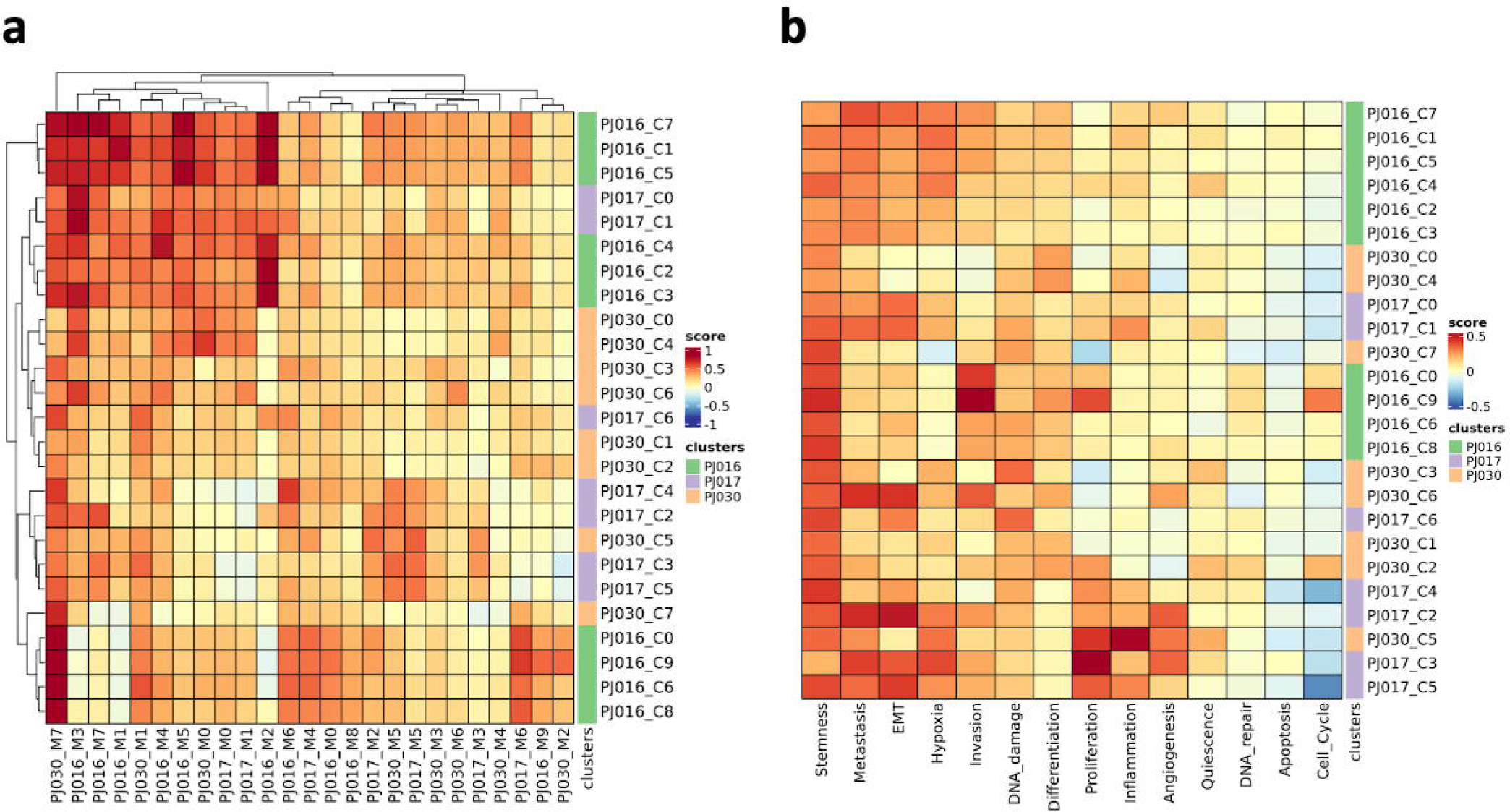
Inter-dataset comparison of gene set expression. **a)** Heatmap of cluster marker (columns, “_Mx” suffix, where “x” indicates cluster number) expression scores across clusters (rows, “_Cx” suffix, where “x” indicates cluster number) of datasets PJ016, PJ017 and PJ030. **b)** Heatmap of CancerSEA functional states scores across clusters of datasets as in panel (a).

The proliferation rate is a relevant indicator for distinguishing cell types in solid tumors and helps to identify cells with potential clinical relevance and interest as candidate therapeutic targets (25,26). Cell proliferation rate is quantified based on the expression of G1/S and G2/M genes:

`scMuffinList <-proliferation_analysis(scMuffinList).`

As a proof-of-concept we jointly analysed cell TC and cell proliferation of sample PJ016 and visualized the results on a diffusion map (27) – which can be run by means of the wrapper function `diff_map()`. This joint analysis shows that cells with the highest values of TC-ratio tend to have lower proliferation scores **(Figure 6d)**. These cells are located more closely to the branching point of the diffusion map **(Figure 6e)**, while the cells with the highest proliferation rates are located at a corner of one of the branches **(Figure 6f)**. This evidence suggests that the cells with high TC and low proliferation are interesting candidates for further analyses aimed at finding TICs in this dataset of HGG. Interestingly, this particular pattern is captured by TC-ratio, but not by TC-LMR and TC-H **(Supplementary Figure S1, Additional File 1)**.

### 3.5 Comparison of datasets

A SC dataset carries an extensive amount of information. The integration of multiple SC datasets is a challenging task and multiple approaches have been proposed to address it (28). Typically, the integrated datasets are computationally demanding due to their huge size. An alternative possibility lies in cross-checking the expression of cluster markers between datasets: the expression of the cluster markers of every dataset is assessed in all considered datasets. For example, Nguyen *et al*. (5) used this approach to study the occurrence of the characteristic cell types of normal mammary gland across samples collected from different subjects.

scMuffin provides a function to quantify the expression of a series of gene sets across multiple datasets. If these gene sets are defined as cluster markers (“cluster_markers_list” in the code below), then the analysis sheds light over the possible presence of cells with similar expression profiles across datasets:

res <-inter_dataset_comparison(seu_obj_list = seu_obj_list, gsl = cluster_markers_list)

We performed this analysis on three datasets (PJ016, PJ017, PJ030) and found cell clusters that are more similar to clusters of other datasets than clusters of the same dataset, suggesting the presence of similar cells **(Figure 8a)**, like cluster 7 of PJ030 (“PJ030_C7”), which is close to clusters {0, 6, 8, 9} of PJ016. Importantly, the same analysis can be done using gene sets with a functional meaning, to obtain a functional characterization of similar cell types across datasets **(Figure 8b)**.

### 3.6 scMuffin and scCancer

scCancer (29) is an R package for automated processing of SC expression data in cancer. The scMUffin and scCancer address the same challenge and are complementary. Indeed, each of them offers some analyses that the other does not **(Supplementary Table S1, Additional File 1)**. Examples of this are cluster association analysis (CSEA and ORA), comparison of multiple cell partitions and inter-dataset gene set expression assessment, available in scMuffin, and cell interaction, expression programs and cancer micro-environment cell classification, provided by scCancer. Other types of analyses are common to both packages but differ by the underlying algorithm and/or implementation. This is the case, for instance, of gene set scoring and CNV analysis.

## 4. Conclusions

Here, we presented scMuffin, an R package that provides a series of functions to perform and integrate various types of analyses on SC expression data. As a proof-of-concept, to illustrate the user interface and its potentialities, we analysed publicly available SC expression datasets of human HGG. We described two examples of integrative analyses which returned particularly interesting findings that would deserve further investigations: the functional characterization of CNVs highlighted a possible link between amplifications of chromosomes 1p and 19p and invasive tumor phenotype; the joint analysis of transcriptional complexity, proliferation rate and cell state trajectories spotted cells that have some characteristics of TICs. While the inference of CNV from SC expression data is a challenging task, subject to various issues (e.g., missing data, number of aberrancies) still it provides insights useful to clarify cell identity, like the distinction between a more aberrant cluster and the others or the distinction of tumoral and normal cells. The function of scMuffin can be combined conveniently to address various tasks. For example, different partitions of the same dataset obtained using the same clustering method with different parameters can be characterized by means of their overlap and gene set expression, to gain insights on the appropriate number of clusters. The analyses offered by scMuffin and the results achieved in this case study show that scMuffin helps addressing the main challenges in the bioinformatics analysis of SC datasets from solid tumors.

## 6. Availability and requirements

**Project name:** scMuffin

**Project home page:** https://github.com/emosca-cnr/scMuffin

**Operating system:** Platform independent

**Programming language:** R (>= 4.0.0)

**Other requirements:** The R Project for Statistical Computing.

**License:** GPL-3

**Any restrictions to use by non-academics:** According to GPL-3

## 7. List of abbreviations

CNV: Copy Number Variation
CSC: cancer stem cells
CSEA: Cell Set Enrichment Analysis
FDR: False discovery rate
ER: Enrichment Ratio
GSEA: Gene Set Enrichment Analysis
GTEx: The Genotype-Tissue Expression project
HGG: high grade glioma
NES: normalized enrichment score
ORA: over representation analysis
SC: single cell
TC-R: Transcriptional Complexity Ratio
TC-LMR: Transcriptional Complexity Linear Model Residuals
TC-H: Transcriptional Complexity Entropy
TICs: Tumor Initiating Cells
UMAP: Uniform Manifold Approximation and Projection

## 8. Declarations

**Ethics approval and consent to participate**. Not applicable.

**Consent for publication**. Not applicable.

**Availability of data and materials**. The data used for the analyses described in this manuscript were obtained from: The Gene Expression Omnibus (11), under the accession GSE103224; the GTEx Portal (17) on 04/08/2020.

## Author’s contributions

VN implemented the CNV functions, carried out the analyses, interpreted the results and wrote the article. AC implemented clustering functions, carried out the analyses, wrote the article, tested the package. NDN drafted the package, carried out the analyses and wrote the article. IC curated the biological aspects of CNV analysis and revised the article. MM and MG set up the computational infrastructure for data analysis. CC and EP curated the biological aspects of solid tumor data analysis. AS tested the package. RR, IZ, LM and AM contributed to the design of the study. PP contributed to the software design, case study definition, interpretation of the results and wrote the article. EM designed the study, implemented the software, performed the analysis, interpreted the results, and wrote the article. All authors read and approved the final manuscript.

## 10 Additional file

Format: PDF

Title: Supplementary material

Description: supplementary methods, table and figure

