## Additional file 1 for "scMuffin: an R package for disentangling solid tumor heterogeneity from single-cell expression data"

**SUPPLEMENTARY METHODS** __ **2**

SINGLE CELL DATA ANALYSIS __ 2

**SUPPLEMENTARY FIGURE** __ **3**

FIGURE S1. TRANSCRIPTIONAL COMPLEXITY, PROLIFERATION RATE AND CELL STATE TRAJECTORIES (PJ016). __ 3

**SUPPLEMENTARY TABLE** __ **4**

TABLE S1. OVERVIEW OF THE ANALYSIS PROVIDED BY SCMUFFIN AND SCCANCER. __ 4

**REFERENCES** __ **5**

### Supplementary Methods

#### Single cell data analysis

The filtered genes-by-cells count matrix of each sample was downloaded from the Gene Expression Omnibus (GEO) repository and was processed using the R package Seurat (1). Cells with less than 1’000 genes and genes expressed in less than 100 cells were excluded. The resulting matrices were log-normalized using the Seurat “NormalizeData” function. In summary, we obtained:

| **Sample (GEO accession)** | **#genes** | **#cells** |
|---|---:|---:|
| PJ016 (GSM2758471) | 12’126 | 2’828 |
| PJ017 (GSM2758472) | 3746 | 642 |
| PJ030 (GSM2758475) | 4’948 | 1’173 |

Cell clustering was performed with Seurat: principal component analysis (PCA) was run on the 2’000 most variable genes, identified by means of “FindVariableFeatures” (method “vst”); “FindNeighbors” was run on the top 10 PCs; “FindClusters” was run with default parameters. The UMAP coordinates were obtained by means of the Seurat “RunUMAP” function, on the top 10 PCs.

### Supplementary figure

#### Figure S1. Transcriptional complexity, proliferation rate and cell state trajectories (PJ016).

**a-c)** Distribution of cells according to the first two “diffusion components” (DC): colors indicate (a) Transcriptional Complexity Linear Model Residual (TC-LMR), (b) Transcriptional Complexity - Entropy (TC-H) and (c) proliferation score. **d-e**) Distribution of (d) TC-LMR and (e) TC-H values split by proliferation score.

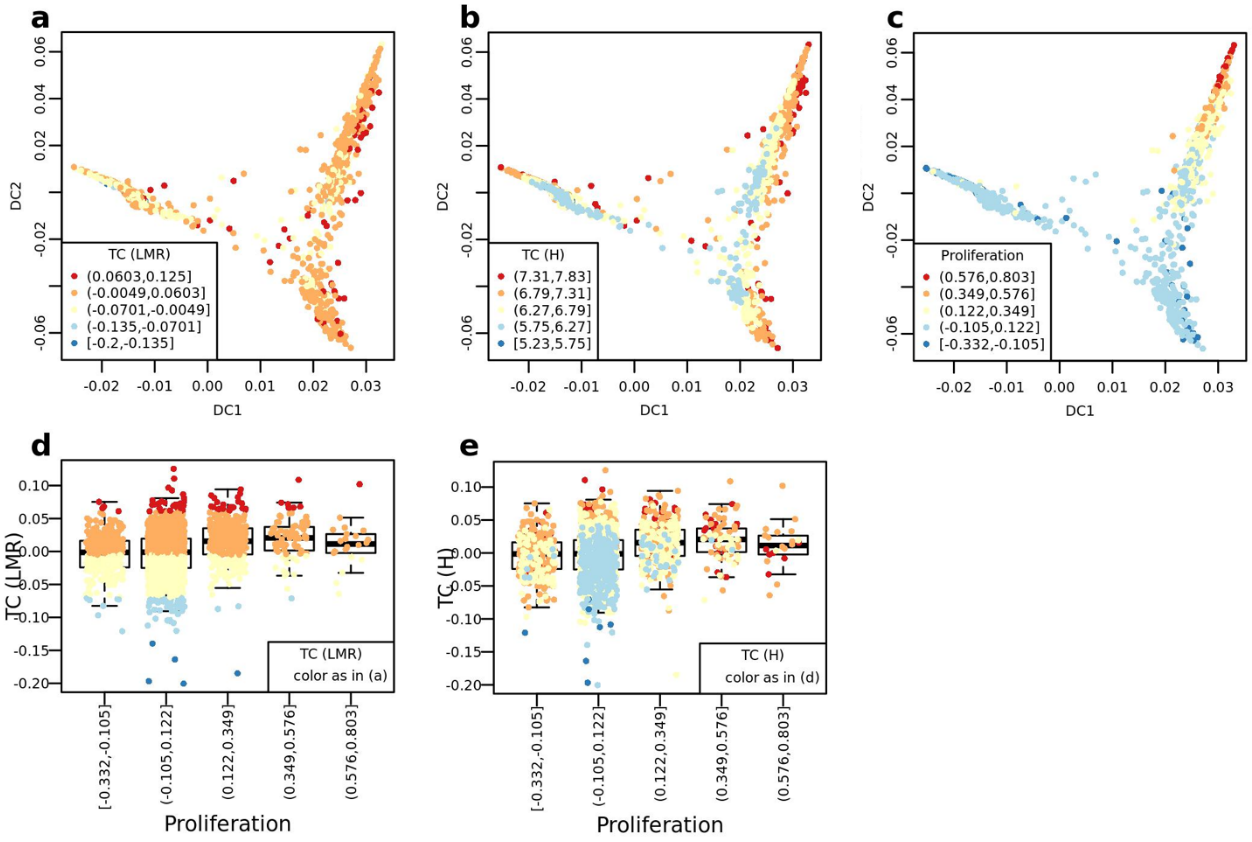

### Supplementary table

#### Table S1. Overview of the analysis provided by scMuffin and scCancer.

| **Analysis** | **scMuffin (this work)** | **scCancer** (2) |
|---|---|---|
| Quality control, filtering, normalization, cell clustering, dimensionality reduction and cluster marker identification | NA | - Cell statistics (nUMI, nGene)<br>- Gene statistics (mitochondrial genes, ribosomal genes, ambient genes)<br>- Based on Seurat (V3 (1)) |
| Cancer micro-environment cell type classification | NA | - data-driven one-class logistic model |
| Gene set scoring | - Algorithm by Tirosh *et al*. (3), with support for missing values<br>- Fine-tuning of parameters<br>- Predefined gene sets: CancerSEA, MSigDB, Cell marker, PanglaoDB<br>- Parallel implementation<br>- Cell- and cluster-level gene set scores<br>- Visualization: Heatmap, UMAP | - GSVA and Seurat AddModuleScore() (based on Tirosh *et al.* (3))<br>- Predefined gene sets: MSigDB Hallmarks<br>- Limited control over parameters<br>- Cell-level gene set scores<br>- Visualization: Heatmap |
| CNV inference | - Based on adjacent gene windows approach by Patel *et al*. (4)<br>- Parallel implementation<br>- CNV region detection<br>- Cell CNV summary score<br>- CNV clusters<br>- Visualizations: CNV Heatmap (with annotation of genes or CNV regions); cluster median CNV profile; cell CNV summary score (per cluster, cell type); | - based on InferCNV algorithm<br>- Cell-level "malignancy" estimation<br>- cell classification (malignant, normal)<br>- Visualization: CNV heatmap, malignancy score over t-SNE, amount of malignant cells per cluster |
| Stemness | - Transcriptional complexity (TR-Ratio, TR-LMR, TR-H) | - Spearman correlation coefficient between cells' expression and stemness signature (One Class Logistic Regression trained over a stem/progenitor database) |
| Comparison of multiple cell partitions | - Support multiple cell partitions<br>- Overlap matrix between all-pairs of clusters | NA |
| Cell cycle/cell proliferation | - Proliferation score (maximum between the gene set scores for G1/S and G2/M) | - gene set scoring (based on Seurat AddModule Score) of cell cycle genes (G2/M and S phase markers) |
| Expression programs | NA | Based on non-negative matrix factorization |
| Cluster association analysis | - quantitative features: Cell Set Enrichment Analysis (CSEA)<br>- categorical features: Over Representation analysis (ORA)<br>- Visualization: boxplots, barplot, heatmap | NA |
| Survival analysis | NA | Survival analysis on gene expression data (patient level) based on marker genes or signatures extracted from SC analysis |
| Cell interaction | NA | Based on FANTOM5 ligand-receptor interaction and algorithm by Kumar *et al.* (5) |

| | | |
|---|---|---|
| Dataset integration | NA | Various algorithms: "NormalMNN". "Harmony", "NormalMNN", "SeuratMNN", "Raw", "Regression" and "LIGER |
| Dataset comparison | Assessment of gene set expression across cell clusters from multiple datasets | NA |
